# The Dynamics of Influenza A H3N2 Defective Viral Genomes from a Human Challenge Study

**DOI:** 10.1101/814673

**Authors:** Michael A. Martin, Drishti Kaul, Gene S. Tan, Christopher W. Woods, Katia Koelle

**Affiliations:** Department of Biology, Emory University, Atlanta, GA, USA; Population Biology, Ecology, and Evolution Graduate Program, Laney Graduate School, Emory University, Atlanta, GA, USA; Infectious Diseases, The J. Craig Venter Institute, La Jolla, California, USA; Division of Infectious Diseases, Department of Medicine, University California San Diego, La Jolla, California, USA; Center for Applied Genomics and Precision Medicine, Duke University School of Medicine, Durham, NC, USA

## Abstract

The rapid evolution of influenza is an important contributing factor to its high worldwide incidence. The emergence and spread of genetic point mutations has been thoroughly studied both within populations and within individual hosts. In addition, influenza viruses are also known to generate genomic variation during their replication in the form of defective viral genomes (DVGs). These DVGs are formed by internal deletions in at least one gene segment that render them incapable of replication without the presence of wild-type virus. DVGs have previously been identified in natural human infections and may be associated with less severe clinical outcomes. These studies have not been able to address how DVG populations evolve *in vivo* in individual infections due to their cross-sectional design. Here we present an analysis of DVGs present in samples from two longitudinal influenza A H3N2 human challenge studies. We observe the generation of DVGs in almost all subjects. Although the genetic composition of DVG populations was highly variable, identical DVGs were observed both between multiple samples within single hosts as well as between hosts. Most likely due to stochastic effects, we did not observe clear instances of selection for specific DVGs or for shorter DVGs over the course of infection. Furthermore, DVG presence was not found to be associated with peak viral titer or peak symptom scores. Our analyses highlight the diversity of DVG populations within a host over the course of infection and the apparent role that genetic drift plays in their population dynamics.

**Importance:** The evolution of influenza virus, in terms of single nucleotide variants and the reassortment of gene segments, has been studied in detail. However, influenza is known to generate defective viral genomes (DVGs) during replication, and little is known about how these genomes evolve both within hosts and at the population level. Studies in animal models have indicated that prophylactically or therapeutically administered DVGs can impact patterns of disease progression. However, the formation of naturally-occurring DVGs, their evolutionary dynamics, and their contribution to disease severity in human hosts is not well understood. Here, we identify the formation of *de novo* DVGs in samples from human challenge studies throughout the course of infection. We analyze their evolutionary trajectories, revealing the important role of genetic drift in shaping DVG populations during acute infections with well-adapted viral strains.

## Introduction

Influenza defective viral genomes (DVGs) were first reported by von Magnus (1) and have since been characterized *in vivo* during high multiplicity of infection (MOI) passage studies (2–7) as well as from clinical human samples (8, 9). DVGs are classified as viral genomes harboring mutations which render them incapable of self-replication. Their propagation depends on replication by wild-type helper virus (10). Influenza DVGs are formed by large internal deletions (11), which retain the 3’ and 5’ untranslated regions that are necessary for replication (12–15) and virion packaging (16–18). Although DVGs have been observed in all eight influenza gene segments (8, 19, 20), they have been most commonly found in the three polymerase genes (PB2, PB1, PA) (19, 21), the longest gene segments of the influenza virus genome.

DVGs that interfere with the replication of wild-type virus have been termed defective interfering particles, or DIPs. It is thought that DIPs are either preferentially replicated (22) and/or packaged (23, 24) given their shorter length. This is thought to lead to the characteristic oscillations in the relative populations of DIP and wild-type virus during passage studies (6) as DIP populations outcompete wild-type virus initially but ultimately crash when the quantity of wild-type virus drops below that necessary to maintain DIP populations. DIPs may also contribute to immune system activation (25, 26). The presence of DVGs during the course of an infection also appears to be associated with less severe clinical outcomes (9). The use of exogenous DIPs has been proposed as a potential therapeutic for influenza, with recent animal studies demonstrating that DIPs administered prophylactically and/or therapeutically can reduce the severity of clinical disease outcomes (27–30).

Influenza A virus (IAV) DVGs have been previously detected in natural human infections from deep sequencing data (8). In this cohort study, DVGs were present in about half of the samples analyzed and were most common in the PB2, PB1, and PA gene segments. A limitation of this study, however, is that it offered only a cross-sectional view of IAV DVG populations. While it has been shown that Sendai virus DVG populations expand during the first 12 hours of infection in a mouse model (25), the evolution of IAV DVG populations within a human host over the course of an infection has not been well characterized.

Here, we report an analysis of IAV DVG populations identified from deep sequencing data taken over the course of infection during two longitudinal human challenge studies with different treatment cohorts. We observe the generation of *de novo* DVGs in nearly all subjects, primarily in the polymerase gene segments (PB2, PB1, and PA). DVG populations were highly variable over time in DVG species composition as well as in DVG species relative abundance. Over the course of infection, individual DVG species were observed to arise, fluctuate in abundance, as well as disappear from the DVG population. Overall, we found no trend towards decreasing diversity of DVG populations or towards shorter DVG species during the five days post challenge, likely due to the dominance of stochastic effects. Furthermore, we were unable to detect an association between DVG levels and peak viral titers, potentially due to the negative feedback between DVG and wild-type virus. Similarly, higher DVG levels were not associated with more severe symptoms. This study helps to illustrate the stochastic dynamics of DVG populations within a host during acute infection with a well-adapted viral strain, a scenario under which fitness variation in the wild-type virus population is expected to be relatively small.

## Materials and Methods

### Ethics statement

The procedures followed in the human challenge studies were in accordance with the Declaration of Helsinki. The studies were approved by the institutional review boards (IRBs) of Duke University Medical Center (Durham, NC), the Space and Naval Warfare Systems Center San Diego (SSD-SD) of the US Department of Defense (Washington, DC), the East London and City Research Ethics Committee 1 (London, UK), and the Independent Western Institutional Review Board (Olympia, WA). All participants provided written consent. *Subject enrollment and challenge study protocol.* Data analyzed in this study were from two previously described human challenge studies (“study 1” indicated by three digit sample IDs and “study 2” indicated by four digit sample IDs beginning with a 5) (31–38). These studies were originally designed to assess changes in host gene expression during the course of influenza infection. Subjects were intranasally inoculated with 3.08 – 6.41 log_10_(TCID_50_/mL) of the challenge virus (“reference strain”). The reference strain was produced by passaging a human isolate of A/Wisconsin/67/2005 (H3N2) [GenBank accession numbers CY114381 to CY114388] three times in avian primary chicken kidney cells, 4 times in embryonated chicken eggs, and twice in GMP Vero cells.

A subset of subjects in study 2 were treated with oseltamivir on the evening of the first day post challenge (“early treatment cohort”). All study one and remaining study two received oseltamivir on the evening of the fifth day post challenge (“standard treatment cohort”). Nasal wash samples were taken at various time-points post-challenge (study 1: 0, 24, 48, 72, 96, 120, 144, 168 hours; study 2: 23, 29, 42, 53, 70, 76.5, 95, 100.5, 118, 124.5, 141.5, 148.5, 165 hours).

Time of peak viral titer was defined as the time from challenge to the earliest time point at which the maximum viral titer was reached. Duration of infection was defined as the time from challenge to the latest positive viral titer.

Modified Jackson symptom scores (41) were also collected throughout the seven days post challenge (study 1: 0, 12, 21, 36, 45, 60, 69, 84, 93, 108, 117, 132, 141, 156, and 164 hours; study 2: 0, 8, 16, 24, 32, 40, 48, 56, 64, 72, 80, 88, 96, 104, 112, 120, 132, 144, 156, 168 hours). Time to peak symptom score was defined as the time from challenge to the earliest time point at which the maximum symptom score was reached. Duration of symptoms was defined as the time from challenge to the last non-zero symptom score or to the end of follow-up, whichever occurred sooner. The association between treatment cohort and clinical data was assessed with Mann-Whitney *U* tests in RStudio v1.1.447 (42).

Previous analyses found no association between inoculum dose and probability of infection. Given infection, inoculum dose was not associated with disease outcome or the amount of viral shedding (34). We thus did not stratify any of our analyses by subject inoculum dose.

### Generation of sequence data

Samples that were IAV positive by cell culture or quantitative PCR were further processed for whole genome sequencing. In brief, the eight genomic RNA segments of IAV were reverse-transcribed and PCR amplified using a multi-segment RT-PCR (39) from whole RNA extracted from nasopharyngeal samples. Individual samples were then barcoded twice using the sequence independent single primer amplification (SISPA) method (40), which involves a primer extension step with a Klenow fragment (37°C for 60 minutes, 75°C for 10 minutes and 4°C hold) [New England Biolabs] and PCR amplification with a DNA polymerase (Preheat at 94°C for 2 minutes followed by 45 cycles of 94°C for 30 seconds, 55°C for 30 seconds and 68°C for 30 seconds with a final extension time of 68°C for 10 minutes and a 4°C hold) [Gotaq, Promega]. To reduce chimerism, PCR products were treated with exonuclease I (37°C for 60 minutes). Separately, a parallel SISPA was performed from the same sample set, but was not treated with exonuclease I. Samples treated with or without exonuclease I were pooled separately and sequenced on an Illumina HiSeq 2000 instrument (Paired-end sequencing; 2 × 100 bp read). SISPA barcoded reads were then demultiplexed and merged based on the barcode sequence, followed by primer and barcode removal and quality trimming using an in-house script at the JCVI. Sequencing runs with or without exonuclease I were used as technical replicates.

### Sequence data analysis

PCR chimeras were removed using a python v2.7 (43) script by identifying forward and reverse reads from the same DNA fragment with conflicting barcodes. Following the removal of chimeric reads, FastQC v0.11.3 (44) was performed on all samples to ensure sequencing quality. Kraken2 v2.0.8-beta (45) with a complete RefSeq viral database was used to identify reads assigned to influenza A, which were then further quality trimmed with Trimmomatic v0.38 (46). Leading or trailing bases with quality < 3 were removed. Reads were cut when the average quality per base in 4-base wide sliding windows was < 15, and reads with less than 50 bases were excluded. Reads were aligned to the reference strain (GenBank CY114381 - CY114388) using STAR v2.7.0e (47). A STAR pre-indexing string of length six was used to generate genome indexing files. SAM files including only uniquely mapped reads were converted to BAM files which were sorted and indexed using SAMtools v1.9 and HTSlib v1.9 (48). Single-nucleotide polymorphisms (SNPs) were called using the BCFtools v1.9 (48) “mpileup,” “call,” and “norm” commands. Only reads with mapping quality ≥ 255 (uniquely mapped) and bases with quality ≥ 20 were used. BCFtools “consensus” was used to generate sample-specific reference genomes including SNPs present in more than 50% of the high-quality reads at a given position. Reads were aligned to sample-specific reference genomes using STAR in basic two pass mode. BAM files including only primary alignments were generated using SAMtools. PCR duplicates were marked and removed using Picard Tools v2.20.02 (49).

Read depth and read length statistics of the final BAM files were calculated using the SAMtools “depth” and “view” commands along with a simple bash script. Read depth and length statistics were combined for both sequencing runs (with and without exonuclease) for the final analysis.

### DVG identification

Split reads (reads with segments mapping to unique locations in the gene implying the presence of large internal deletions) were identified using a Python v2.7 script and pysam 0.15.2 (https://github.com/pysam-developers/pysam). Split reads with at least 15 alignment matches to the reference, a minimum of five consecutive alignment reference matches, no more than three small indels, and a minimum of 100 consecutive deleted reference bases were used to generate a filtered BAM file. Junction sites for individual DVGs were identified from the “jI” SAM tag, tabulated, and normalized to the total number of reads aligned to that gene segment (norm. DVG reads) using a bash script. Split read depth was calculated using the SAMtools “depth” command.

In order to reduce the number of spurious DVGs, we included only DVGs on a per sample-day basis which were identified in both sequencing runs (with and without exonuclease I) in the final analysis. Raw and normalized DVG read support measurements were combined between the technical replicates. All bioinformatic analyses were performed at the Pittsburgh Supercomputing Center using the Bridges resources.

We define a “DVG species” as DVGs with identical deletion breakpoints and “DVG populations” as all of the observed DVG species within a sample. DVG species are identified by the first and last reference bases deleted (first_last). DVG load for a given gene segment is defined as the sum of the normalized DVG read count for all observed DVGs. DVG load for all gene segments is the average of the normalized DVG read count over the eight genes. The effect of treatment cohort on peak DVG load was assessed using a Mann-Whitney *U* test calculated in RStudio. The association between DVG presence on different gene segments was assessed using Fisher’s exact test calculated in RStudio.

### DVG diversity calculation

To evaluate the degree of DVG diversity within a sample while adjusting for the variable number of observed DVG species, we utilized Pielou’s evenness index (50), given by 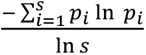 where *s* is the number of DVG species and *p_i_* is the proportion of DVG reads which support that DVG species. An evenness of 1 corresponds to a population in which all observed species are present at the same frequency. This metric was calculated in RStudio.

All figures were generated in RStudio using ggplot2 v3.1.0 (51) and cowplot v0.9.3 (52).

Raw sequencing data are accessible under NCBI BioProject PRJNA577644. Scripts used for the generation of data and figures in this report are available at https://github.com/koellelab/IAV_human_challenge_study_code.

## Results

### Data summary

Of the 37 participants in the human challenge studies, 17 were successfully infected and had at least one sample successfully sequenced. Seven of these 17 individuals belonged to the early treatment cohort; the remaining ten belonged to the standard treatment cohort. Peak viral titers ranged from 1.75 to 6.25 log_10_(TCID_50_/mL) (mean [population standard deviation (sd)]: 4.5 [1.2] log_10_(TCID_50_/mL)) and occurred 24 to 120 hours post-challenge (mean [sd]: 58 [26] hours)). Peak viral titer did not appear to differ between the early and standard treatment group (mean [sd]: 4.3 [1.5] v. 4.6 [0.8] log_10_(TCID_50_/mL); p-value = 0.922). However, those in the early treatment cohort tended to reach peak viral titer faster (cohort (mean [sd]: 48 [20] v. 65 [27] hours; p-value = 0.080) and tended to have shorter durations of infection (mean [sd]: 74 [22] v. 118 [37] hours; p-value = 0.035; Figure S1a, Table S1, Table S2).

Peak total symptom scores ranged from 1 to 16 (mean [sd]: 6.5 [4.8]) and did not differ significantly between treatment cohorts (p-value = 0.922). Time to peak symptom score was shorter in the early treatment cohort (mean [sd]: 29 [14] v. 59 [18]; p-value = 0.003) as was the duration of symptoms (mean [sd]: 83 [37] v. 134 [17]; p-value: 0.005). Cumulative symptom scores were highly variable between subjects and no difference was observed between the two cohorts (mean [sd]: 43 [50] v. 35 [29]; p-value = 0.922; Figure S1b, Table S1, Table S2).

A total of 43 samples, including the inoculum, were successfully deep-sequenced. The number of successfully sequenced samples per subject ranged from one to five (Figure 1A). Following read trimming, the per-sample, per-gene average read length ranged from 70 to 72 nucleotides (nt). The average genome-wide read depth was 118 reads (range: 63 to 166) (Table S3, Figure S2A, Figure S2B).

**Figure 1.**
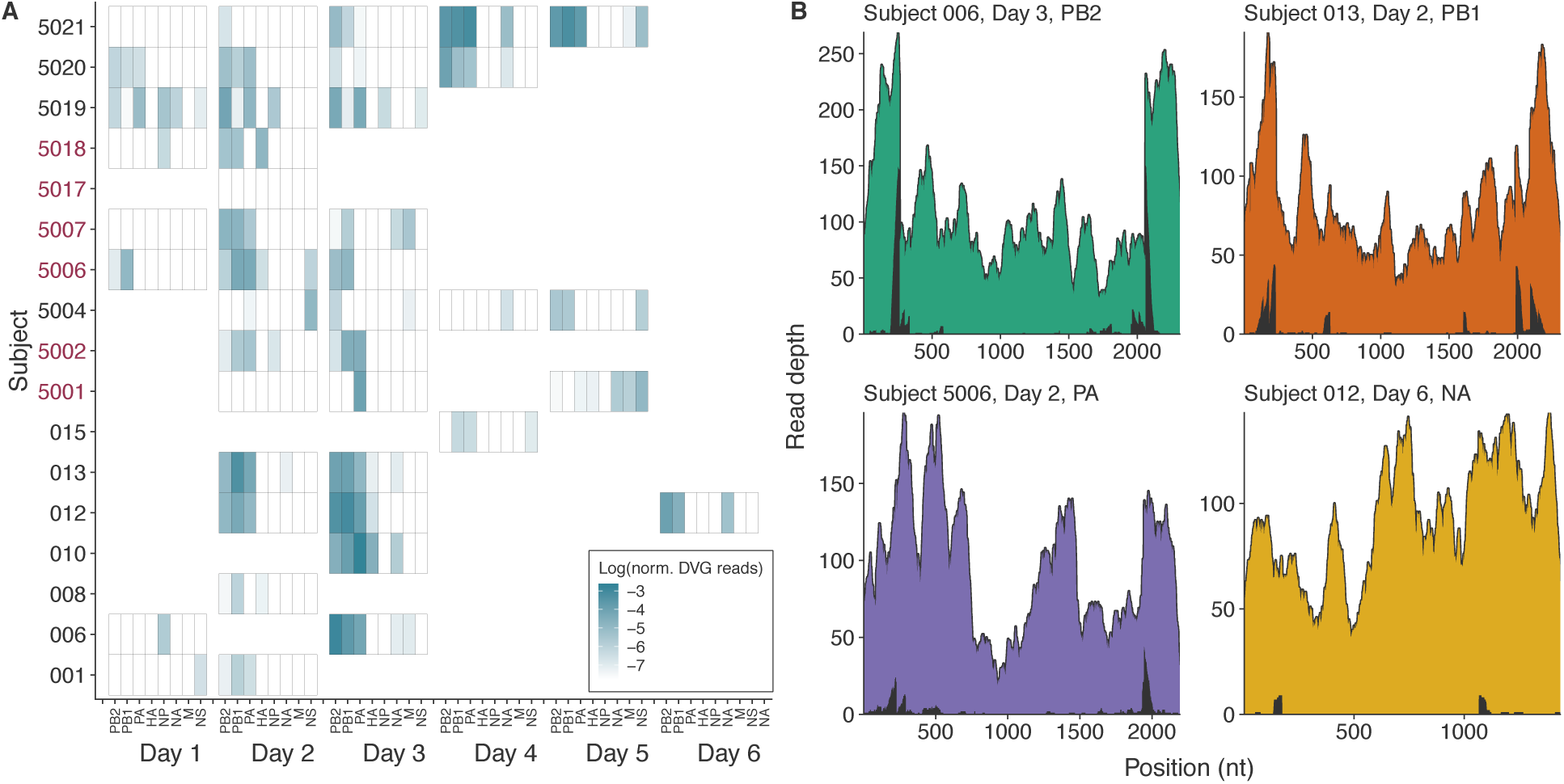
Graphical summary of the data used in this study. A) Heatmap showing the number of sequencing reads indicating the presence of defective viral genomes in each gene segment normalized by the total number of sequencing reads aligned to that gene segment. Rows represent individual subjects (red text indicates early treatment (oseltamivir on the evening of the first day post challenge) cohort). Columns represent day of sampling; sub columns indicate gene segment. White space indicates lack of sequencing data. B) Representative coverage plots in various gene segments. Background colored area shows the total read depth at a given nucleotide (nt) position. Black color in the foreground represents the coverage depth of split reads, indicative of DVGs.

### Widely observed de novo DVGs

DVGs in the challenge stock were largely absent, identified at only low levels in the NP, NA, and NS gene segments and absent from the other gene segments (Figure S3). On the contrary, DVGs were observed in all but one successfully infected subject (Figure 1). For the subject in which no DVGs were detected (subject 5017), only a single sample (day two) was sequenced. This subject was in the early treatment cohort and was also positive for influenza on day three, however, this sample was not successfully sequenced.

Amongst the other subjects, we observed DVGs in at least one of IAV’s eight gene segments in 38/41 successfully sequenced samples. As expected, DVGs were more commonly observed in the polymerase genes (PB2 (n = 31), PB1 (n = 28), and PA (n = 26) v. HA (n=7), NP (n = 7), NA (n = 12), M (5), and NS (10)). DVGs were observed as early as day one and as late as day six post challenge. We did not observe a difference in peak DVG load between treatment cohorts (p-value: 0.713).

Normalized DVG read counts, a proxy for the number of DVGs relative to wild-type virus *in vivo*, ranged from 0.0004 to 0.064 (Table S4, Figure S2C). Among samples with DVGs, deletions in polymerase genes tended to have higher normalized DVG counts, indicating the presence of more DVGs generated from these segments within a given host. Presence of DVGs in the PB2, PB1, and PA gene segments were positively associated with one another (PB2 × PB1 p-value: 2.5 × 10^-6^; PB2 x PA p-value: 5.2 × 10^-3^; PB1 x PA p-value: 2.5 × 10^-3^; Figure S4).

To determine whether certain junction locations were favored in identified DVG species, we tabulated the most commonly observed 3’ and 5’ junction sites (Figure S5, Table S5). Unsurprisingly, most 3’ junction locations were located in the first 500 nt of each gene and the 5’ junction locations in the last 500 nt of each gene. We observed no junction locations within 40 nt of either end of the three polymerase genes, consistent with the theory that the sequences at either end are necessary for replication (12–15) and virion packaging (16–18). The mean [sd] (weighted by normalized DVG read support) number of deleted nucleotides was comparable between gene segments (PB2: 1646 [392]; PB1: 1625 [397], PA: 1593 [332]). A small number of DVGs were observed with 3’ junction sites located towards the 5’ end of a given gene segment. With the exception of a single PB2 DVG, these tended to be found in a small number of samples with low normalized DVG read support. However, 1482_2101 in PB2 was observed in 0.0045 and 0.0068 normalized reads in subject 5020 on days two and four, respectively. This was the dominant DVG present at day two and one of a number of codominant DVGs present at day four.

We observed the presence of identical DVG species across both samples and subjects in the PB2 (n = 16), PB1 (n = 13), PA (n = 13), NP (n = 2), NA (n = 2), and NS (n = 1) genes (Table S6). Repeat DVGs were most commonly present in multiple samples from the same subject (n = 37). For example, the PB2 deletion from nt 356 to nt 1937 in the reference sequence was observed in subject 5021 at four consecutive time points (day two through day five). However, a number were also present in multiple subjects (n = 10). For example, DVG 476_703 in the PB2 gene segment was observed in subject 5006 at day one, subjects 5019 and 5020 at day 2, and subject 5021 at day three. DVG 696_1378 in the NP gene segment was observed in the challenge stock as well as in subject 5002 (day two) and subject 5019 (day two and three, but not one). Given its identification in only these 4 samples, it is unclear whether this DVG was transmitted during challenge or whether it appeared *de novo* in these two subjects (with lack of detection on day one in subject 5019).

### Dynamic within-host DVG populations

Due to the longitudinal nature of these data, we wished to analyze whether systematic changes in the DVG populations within individual hosts were evident. Specifically, we looked at the population composition within hosts to determine whether there was evidence of positive selection for specific DVG species or for changes in the composite characteristics of DVG populations. However, given the acute nature of the infections in this study, any positive selection may be overwhelmed by stochastic effects, as has been previously described for point mutations in acute influenza infections (53).

Our analyses revealed that the composition of DVG populations changed rapidly within-hosts. Individual DVG species were found to rise and fall in their relative read support, and DVG species arose and disappeared throughout the course of infection (Figure S6). For example, in subject 013 the number of individual DVG species increased notably between days two and three, more than doubling in the PB2 and PB1 genes (Figure 2A). However, in general the DVGs observed at day two were still present in the sample at day three. Considerably different dynamics were observed in subject 5021 (Figure S6), from which we have data for day one through four. DVG populations in this subject underwent considerable turnover on day three. Amongst the two PB2 DVGs observed at day two, neither were observed at day three. However, one of these two was observed again at day four. Similar dynamics were observed in both the PB1 gene (in which no DVGs were observed at day three) and the PA gene (in which a unique DVG was observed only at day three).

**Figure 2.**
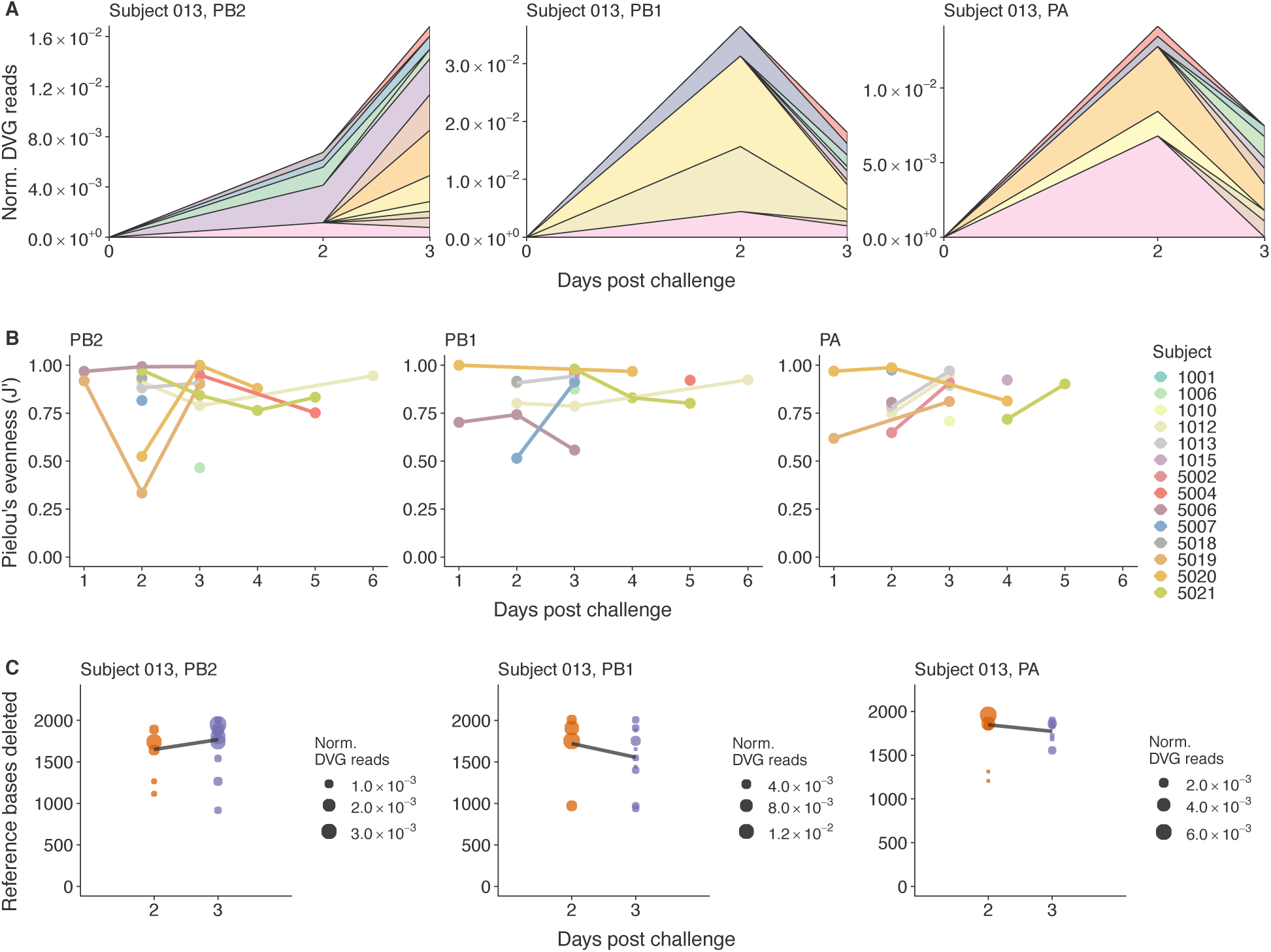
Population dynamics of observed defective viral genome (DVG) populations. A) Stacked area plots showing the DVG species observed at two and three days post challenge in subject 013 in the PB2, PB1, and PA gene segments. Each color represents an individual DVG species. The height of each region represents the normalized number of DVG reads supporting that DVG. B) Diversity of DVG populations in the PB2, PB1, and PA gene segments for each subject in the study. Lines connect data points from the same subject at multiple time points. Diversity is measured my Pielou’s evenness (J’), which is given by 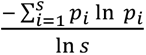 where s is the number of DVG species and *p_i_* is the proportion of DVG reads which support that DVG species. C) Distribution of the number of deleted references bases in each observed DVG species in the PB2, PB1, and PA gene segments of subjects 013. Dot size represents the normalized number of DVG reads supporting a specific DVG species. Trend lines connect the mean number of reference bases deleted at each day, weighted by the normalized DVG read support.

The generation of novel DVG species between sampling timepoints was very common. However, in most cases at any given time point a small number of DVG species accounted for the majority of the relative read support. In certain cases, these dominant DVG species were consistent between time points, however in others the dominant DVG species varied considerably between time points. This suggests that while many different DVG species can be formed during viral replication, stochastic effects during DVG generation or a selective advantage of certain DVG species leads to the observed DVG species unevenness within hosts at any given time point.

To quantitatively assess whether there was evidence for selection for specific DVG species, we determined the trajectory of DVG diversity, measured by Pielou’s evenness (J’), over the course of infection in individual subjects (Figure 2B). We did not observe a trend towards decreasing diversity over the course of infection. For example, amongst PB2 DVGs, subject 5004 as well as subject 5021 witnessed net decreases in DVG diversity overtime whereas there was limited change in J’ for subjects 012 and 5006. Subject 5019 experienced only a transient reduction in diversity on day two when the DVG population was dominated by a single DVG (172_2079). In contrast, subject 5020 experienced a transient increase in diversity on days two and three. Similar stochastic patterns of diversity were also observed in PB1 and PA DVGs.

It has been proposed that DVG species are preferentially replicated over wild-type virus due to their shorter length (22). Therefore, it is reasonable to hypothesize that selection might lead to the evolution of shorter DVG species over the course of infection. To address this hypothesis, we analyzed the number of reference bases deleted over the course of infection amongst those subjects with DVGs in a specific gene at multiple time points (Figure S7). Our data indicated no systematic change in DVG length over the course of infection (Figure 2C, Figure S8). In certain subjects, DVGs tended to get shorter (subject 5004 PB2, subject 5019 PB1, subject 5020 PB2), however in others there was no discernible change in DVG length (subject 012 PB2 and PB1, subject 5002 PA). These results suggest that either genetic drift overwhelms selective forces or that factors other than DVG length more strongly affect DVG fitness.

### Correlation between DVG presence and clinical data

While this study was not powered to detect statistically significant associations between the presence of DVGs and clinical data, we wished to see if there were any qualitative correlations. We first analyzed the relationship between peak viral titer and the peak DVG load within a subject (Figure 3A), expecting a positive correlation as wild-type virus is necessary for the replication of DVGs. We were unable to detect an association between the two measures in this study. The cyclical nature of relative DVG abundance is well established *in vitro* (6) and it is possible that while wild-type virus is necessary for DVG replication, the inhibitory effect of DVG replication on the amount of wild-type virus which is replicated (thereby reducing peak viral titers) obscures any obvious correlation between the two.

**Figure 3.**
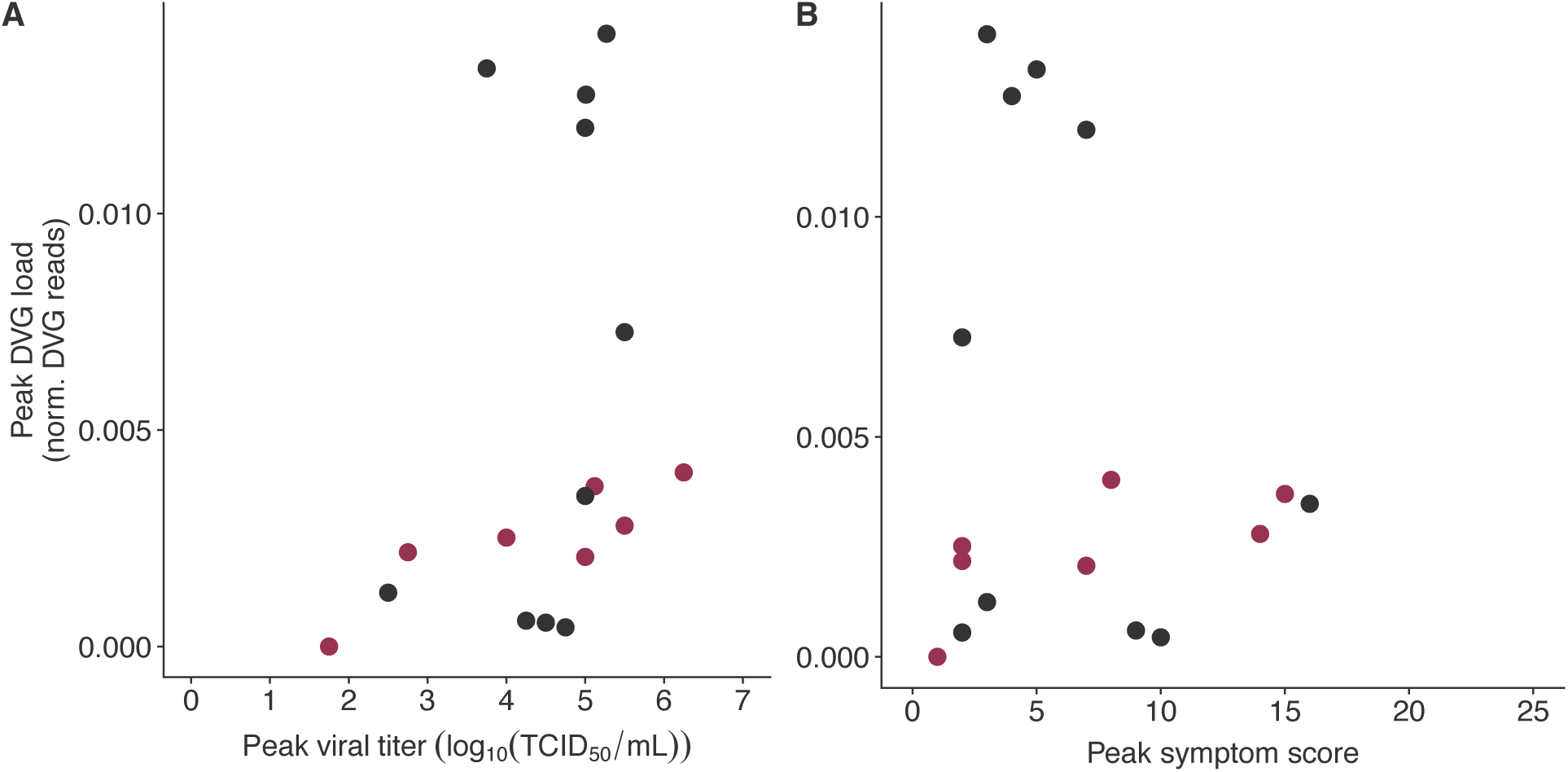
Dot plots of clinical data against the amount of defective viral genomes (DVGs). Each dot represents a subject. A) Dot plot showing the lack of association between peak viral titer (log10(TCID50/mL) on the x-axis and the peak number of normalized DVG reads on the y-axis. B) Dot plot showing the lack of association between peak Modified Jackson symptom score on the x-axis and the peak number of normalized DVG reads on the y-axis.

Previous reports have suggested an associated between DVG presence and IAV clinical outcomes (9). While none of the subjects in this controlled challenge study experienced severe clinical outcomes, we analyzed the association between self-reported symptom scores and DVG presence. As DVGs are thought to dampen the clinical manifestation of IAV infection, we expected high DVG levels to be associated with less severe symptom scores. However, we were unable to detect an association between peak DVG load and peak symptom score in this study (Figure 3B). This lack of association may be due to the contrasting effects of DVGs interfering with wild-type virus replication and packaging and their activation of the innate immune response (25, 26), which is known to be responsible for influenza symptom manifestation. Furthermore, our inability to find an association between DVG load and symptoms may be because the seasonal IAV strain used in this study is relatively avirulent and all subjects were healthy, leading to relatively mild clinical presentations.

## Discussion

The presence of defective influenza genomes has been well characterized in cell cultures (2–7) and animal models (25). Influenza DVGs have also been observed in clinical human H1N1 samples (8). However, to date, no studies have performed longitudinal analyses of naturally occurring DVG populations within humans. Here we presented an analysis of DVG populations in samples from two H3N2 human challenge studies with different treatment protocols for up to seven days post-challenge.

Our analysis supports prior findings that DVG presence is nearly universal and most commonly found on the polymerase gene segments (8), both in terms of presence/absence as well as abundance. Furthermore, we observed that DVGs are often found on multiple polymerase genes within the same subject. The timing of oseltamivir treatment was not found to affect the peak viral titer, peak symptom score, or cumulative symptom scores, nor did it effect the accumulation of DVGs within a host. We observed identical DVG species within single hosts at multiple time points as well as across multiple hosts. As identical DVG species were more likely to be observed within, as opposed to between hosts, this implies that ongoing within-host replication of DVG species following their stochastic generation is likely driving this phenomenon.

DVG populations were shown to be to be highly dynamic in terms of both the DVG species they comprised and the abundance levels of these species. There was no evidence for decreased diversity of DVG populations within a host over the course of infection. Furthermore, we saw no trend towards DVG species becoming on average shorter over time. These results imply that *in vivo* genetic drift may be overwhelming selective forces in shaping the evolutionary dynamics of DVG species in this study. IAV genetic drift playing a strong role in these human challenge studies is not unanticipated, given that the challenge reference strain was a seasonal influenza strain that was relatively well adapted to human hosts and that egg- and cell culture-adapted variants were quickly excluded from the *in vivo* viral populations (31). The effect of spatial structure within the host respiratory system (54) may further augment the effects of genetic drift on DVG populations.

We did not observe an association between peak viral titer or peak symptom score and peak DVG loads. This points to the complex feedback mechanisms which govern the amount of DVG and wild-type virus within a host as well as between the replicative inhibitory effect of DVG generation on wild-type virus replication and the interaction between DVGs and the host immune response.

This analysis has several limitations, largely due to the nature of the available data. The sequencing reads were generated in 2013 and therefore read lengths are shorter and mean read coverage is lower than in more recently generated viral deep sequencing datasets. Furthermore, we did not confirm the presence of DVG species using PCR, as has been done in other studies (8) because samples from these human challenge studies are no longer available. Furthermore, a certain level of noise in the bioinformatics pipeline used to identify DVGs is to be expected. In order to reduce this noise we analyzed only DVG species present in both technical replicates (with and without exonuclease I), however, we opted not to remove DVG species with low supporting read counts as has been previously proposed (55) in order to maintain sensitivity in our measure of DVG diversity. The very low number of split reads observed in the non-polymerase genes, which are known to rarely form DVGs, implies that the level of noise in our analysis is relatively low.

Furthermore, sequencing data were only available at most once per day for each subject and therefore we were unable to assess the fine-scale evolution of DVG species. This sparse sampling is likely why observed DVG populations were so variable between time points. With more frequent sampling we predict it would be possible to observe more gradual transitions between DVG population compositions within a host.

Despite these limitations, this study adds to the growing body of evidence that influenza DVGs are present during human infections and evolve over the course of infection. While the expansion of DVGs in individual infections are surely impacted by wild-type viral dynamics, whether DVGs in turn play a role in shaping infection dynamics and determining disease progression remains an open question.

Further studies with greater temporal resolution and sequencing to a higher read depth may help to more precisely characterize the evolutionary trajectory of DVG populations within individual hosts. Analysis of the most common DVG species observed in future studies may reveal factors that impact DVG stability over the course of an infection. A thorough understanding of the interaction between wild-type virus, DVGs, and the host immune response may ultimately aid in the development of therapeutics based on exogeneous DIPs.

## Acknowledgements

We thank Ashley Sobel Leonard for guidance in accessing and interpreting the available sequence data; Katherine E. E. Johnson, Tim Song, and Elodie Ghedin for explaining their DVG identification pipeline, which was adapted for this analysis; Fadi G. Alnaji and Christopher B. Brooke for their advice on the identification of DVGs from Illumina sequencing data; and David E. Wentworth for his assistance in interpreting the JCVI sequencing methods used to generate the data analyzed here. This work used the Extreme Science and Engineering Discovery Environment (XSEDE) Bridges Regular Memory resource at the Pittsburgh Supercomputing Center (56) and was funded by the US Defense Advanced Research Projects Agency (DARPA) INTERCEPT W911NF-17-2-0034 contract. The funding for the challenge studies was provided by DARPA contract NC66001-07-C-2024. Work was partially funded by the National Institute of Allergy and Infectious Diseases, National Institute of Health award numbers HHSN272200900007C and U19AI110819).

## Supplemental Material

**Figure S1.**
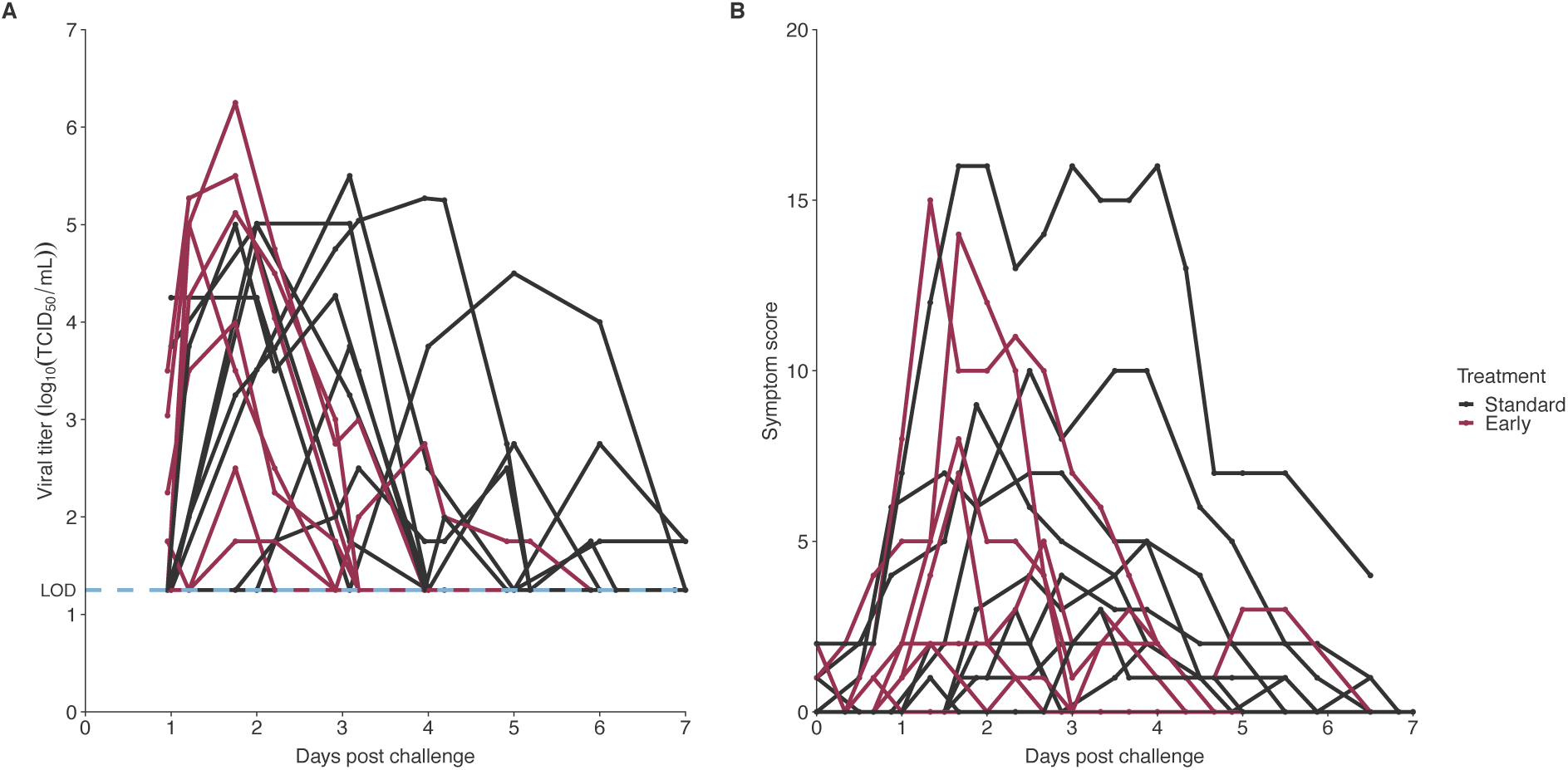
Clinical data included in the study. Red lines represent subjects in the early treatment (oseltamivir on the evening of the first day post challenge) cohort, grey lines represent subjects in the standard treatment (oseltamivir on the evening of the fifth day post challenge) cohort. A) Viral titer (log_10_(TCID_50_/mL)) measurements for each subject at various time-points post-challenge. Blue dotted line at 1.25 log_10_(TCID_50_/mL) represents the limit of detection of the assay used. B) Modified Jackson symptom scores for each subject at various time-points post-challenge.

**Figure S2.**
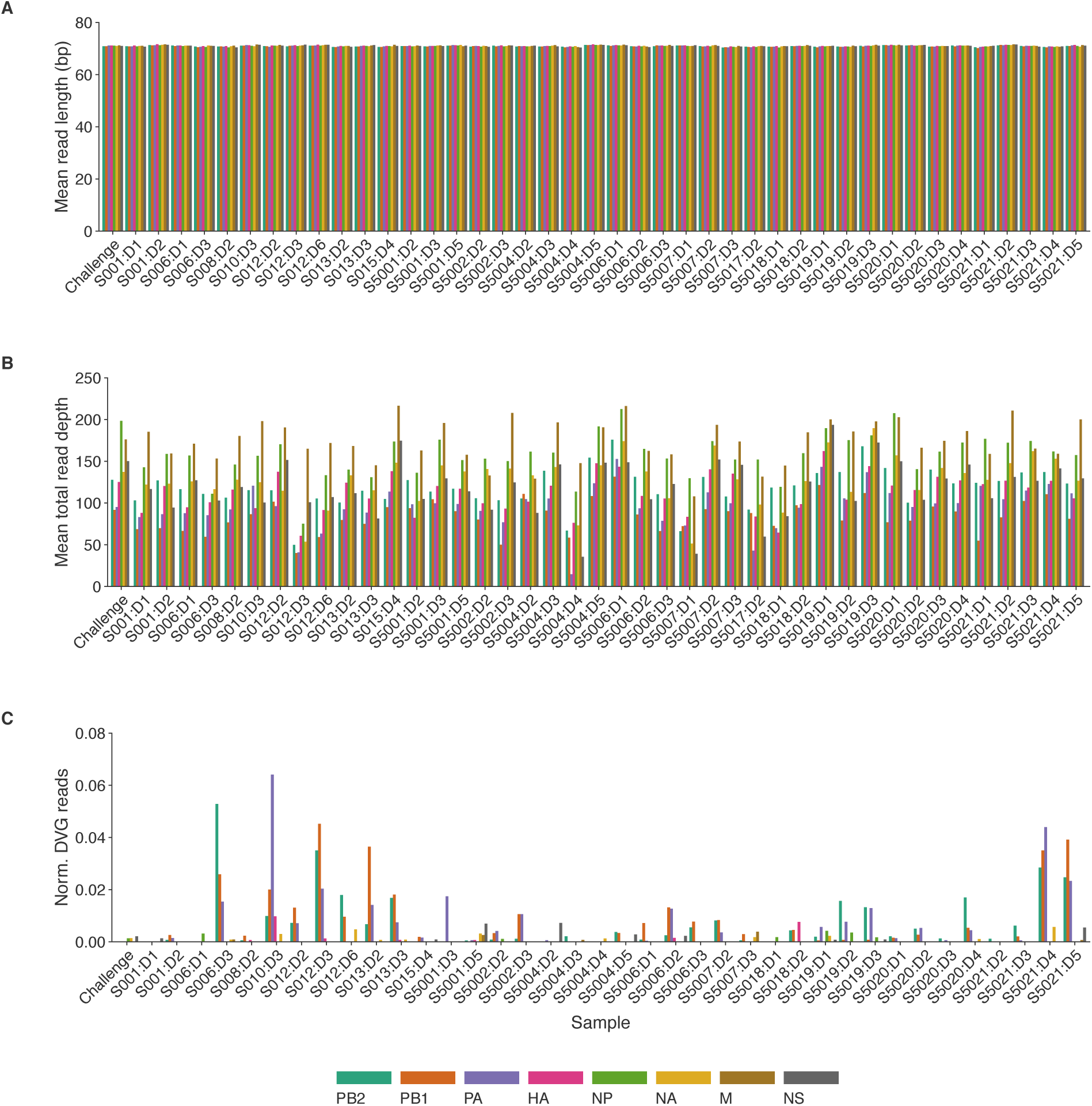
Summary of sequencing data. Bars are organized first by subject, then by sampling day, and finally by gene. Genes are also represented by colors. Data are summed across technical replicates. A) Mean length of mapped reads following read trimming, reference mapping, and removal of PCR duplicates. B) Mean total read depth following read trimming, reference mapping, and removal of PCR duplicates. C) Mean depth of split reads following read trimming, reference mapping, removal of PCR duplicates, and filtering of split reads to exclude those with less than 15 alignment matches, less than 5 consecutive alignment matches, more than three small indels, and with junction locations less than 100 bases apart.

**Figure S3.**
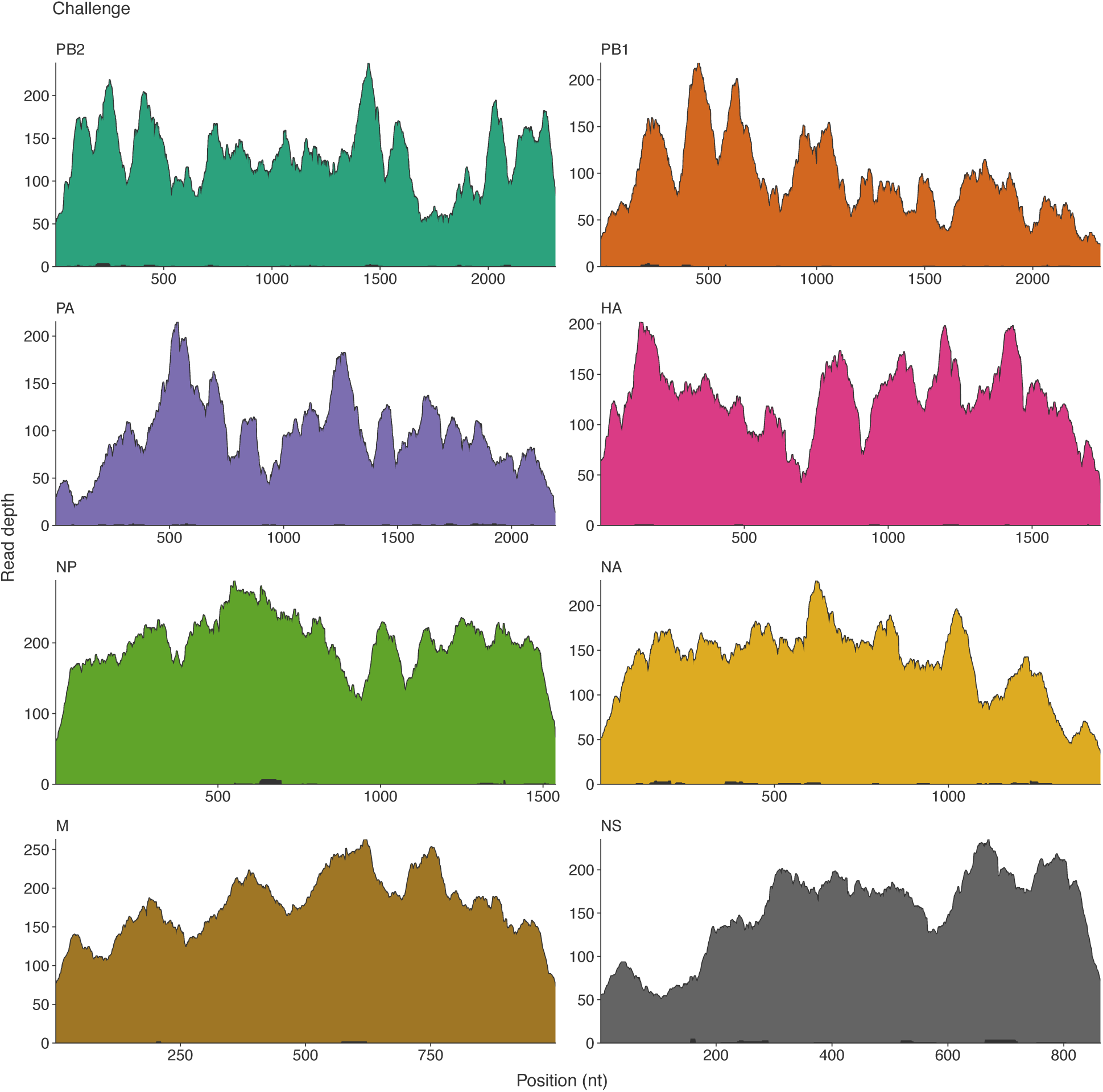
Coverage plot for each gene segment in the challenge stock used to infect all patients. Colored portions represent total read depth and black overlays represent read depth of split reads. Lack of appreciable split read depth indicates a lack of defective viral genomes (DVGs). The NP, NA, and NS gene segments each harbored a single DVG species, while no DVG species were identified in the other 5 gene segments.

**Figure S4.**
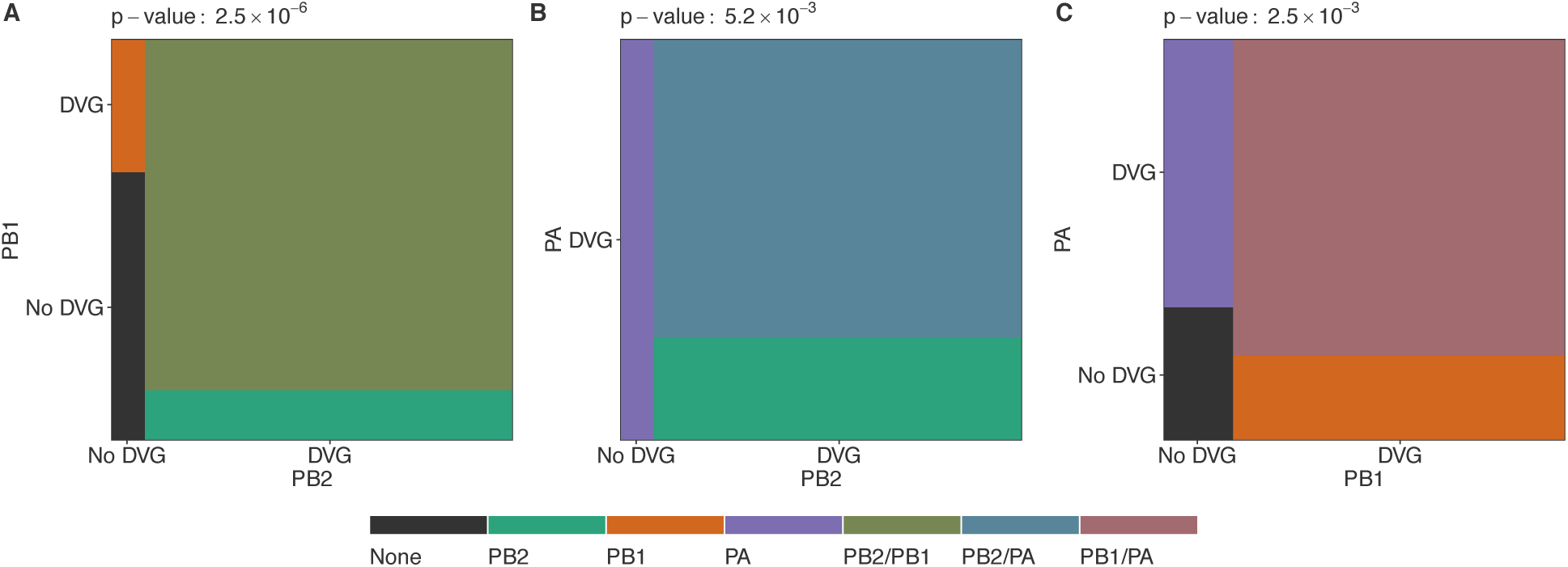
Mosaic plots representing the per-sample dependence of defective viral genome (DVG) presence in pairwise combinations of the PB2, PB1, and PA gene segments. Axis represent the proportion of all samples. Grey area indicates lack of DVGs in either gene segment and colored areas represent DVG presence in either or both gene segments. Fisher’s exact test p-values are listed above each plot. A) Association between DVG presence in the PB2 and PB1 genes. Green area represents samples with DVGs only in the PB2 gene, orange area represents samples with DVGs only in the PB1 gene and forest green area represents samples with DVGs in both genes. B) Association between DVG presence in the PB2 and PA gene segments. Green area represents samples with DVGs only in the PB2 gene, purple area represents samples with DVGs only in the PA gene, and blue area represents samples with DVGs in both. C) Association between DVG presence in the PB1 and PA gene segments. Orange area represents samples with DVGs only in the PB1 gene, purple area represents samples with DVGs only in the PA gene, and mauve area represents samples with DVGs in both genes.

**Figure S5.**
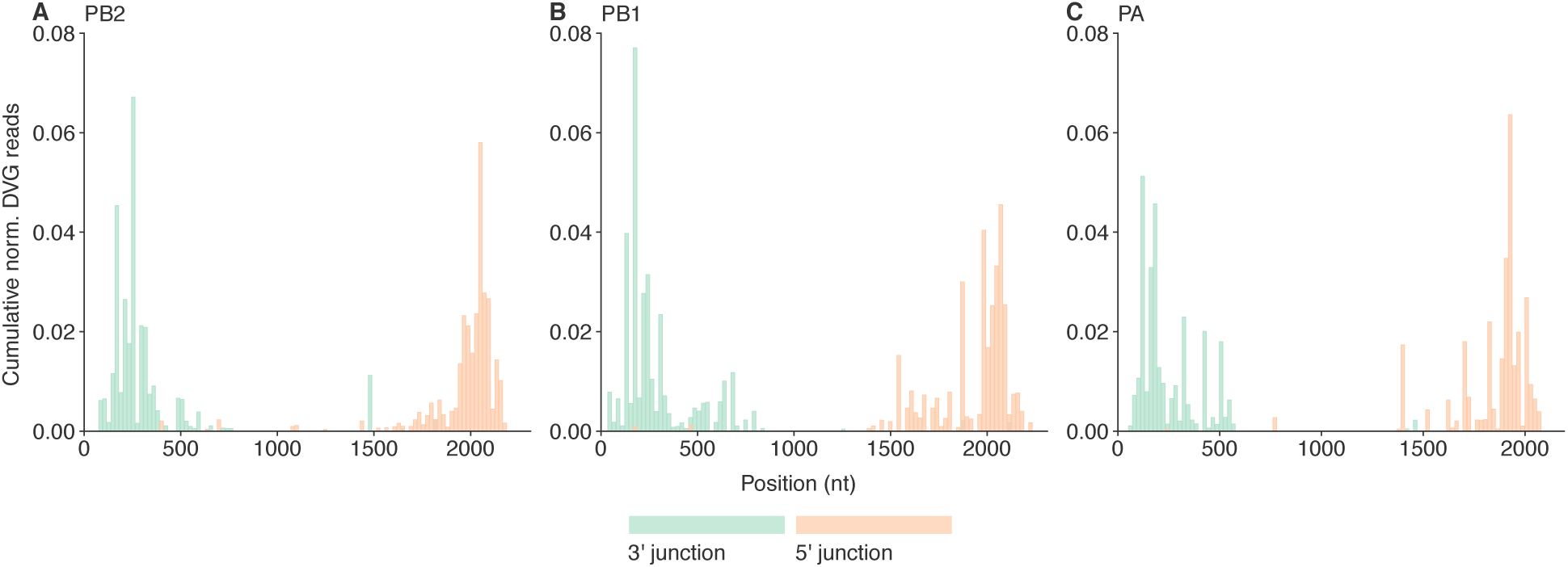
Histograms (100 bins) showing the cumulative relative read support values for the 3’ (green) and 5’ (orange) junction locations (compared to the reference) of the deletions generating individual DVG species. The relative read support for DVGs observed within individual samples was first normalized to the total number of reads mapped to that gene segment and then summed across subjects. Data are shown for DVGs observed in the A) PB2, B) PB1, and C) PA genes.

**Figure S6.**
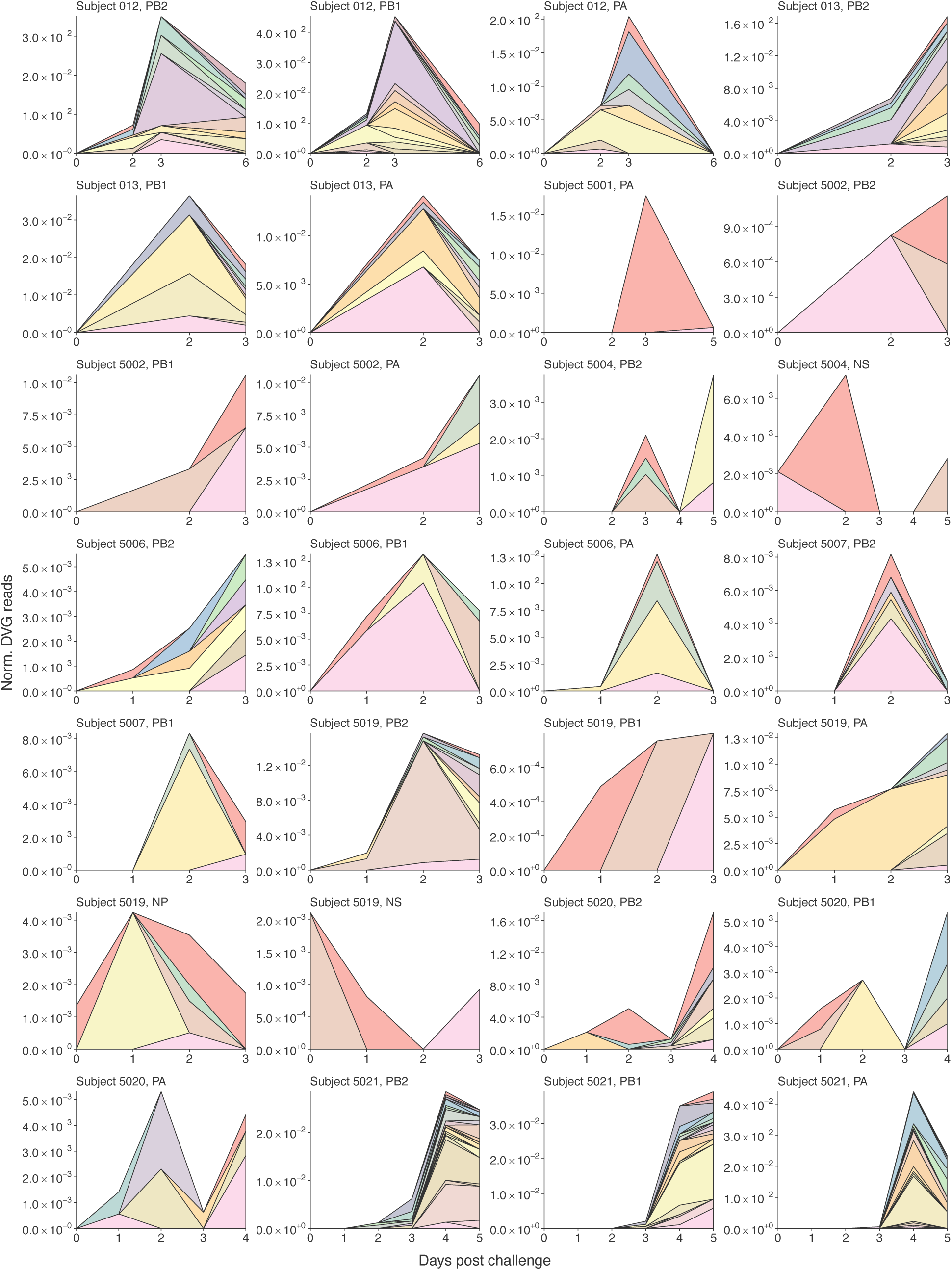
Stacked area plots representing the defective viral genome (DVG) species within each gene for subjects with DVGs observed on multiple days. Each color represents an individual DVG species. The height of each region represents the normalized number of DVG reads supporting that DVG. DVG colors are not consistent between subjects.

**Figure S7.**
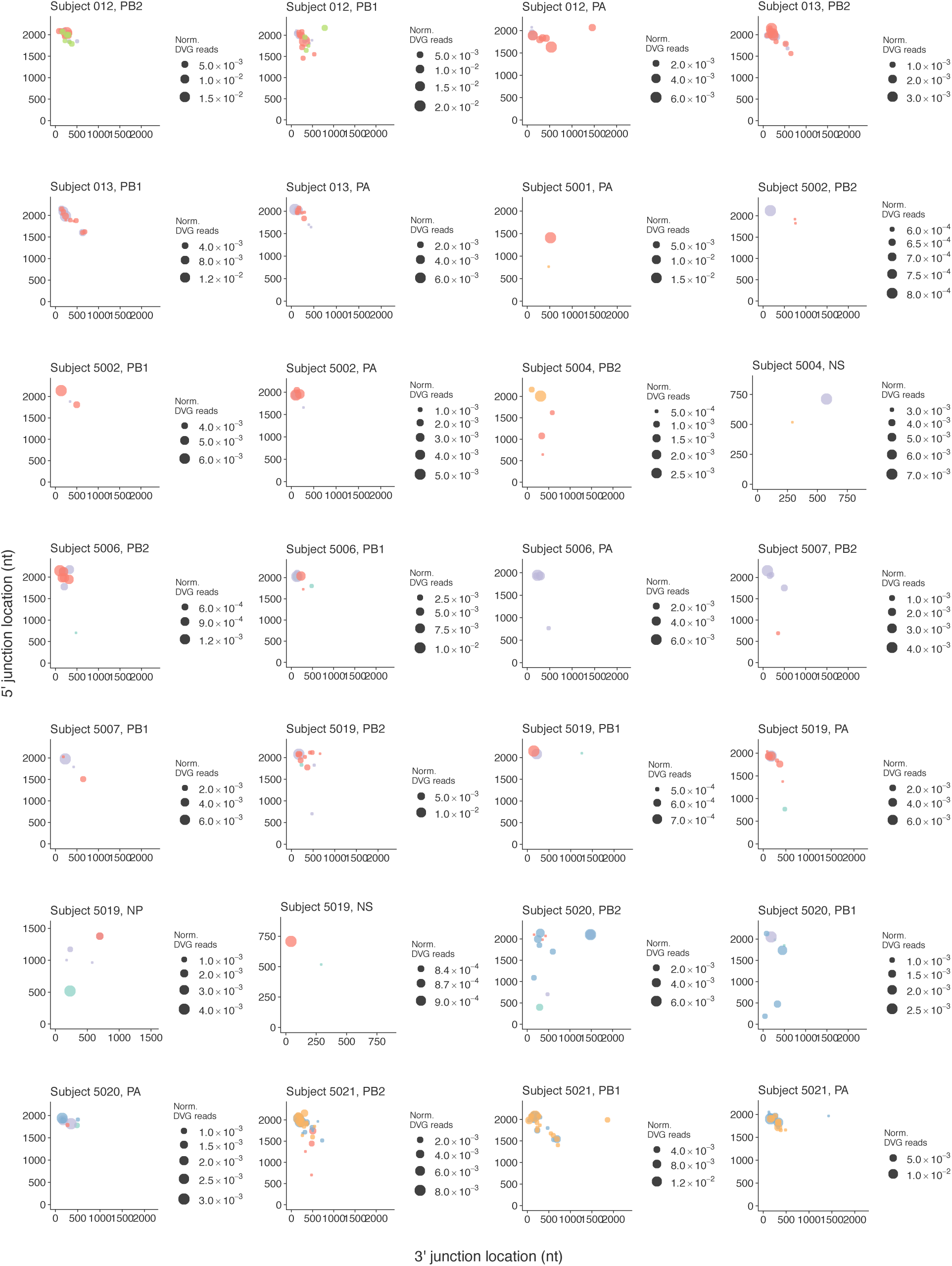
Dot plots representing the defective viral genome (DVG) species within each gene for subjects with DVGs observed on multiple days. The x-axis represents the 3’ junction location and the y-axis represents the 5’ junction location. Dot size is dependent on the number of reads supporting a given DVG species, normalized by the number of reads mapped to that gene segment. Color represents sampling day.

**Figure S8.**
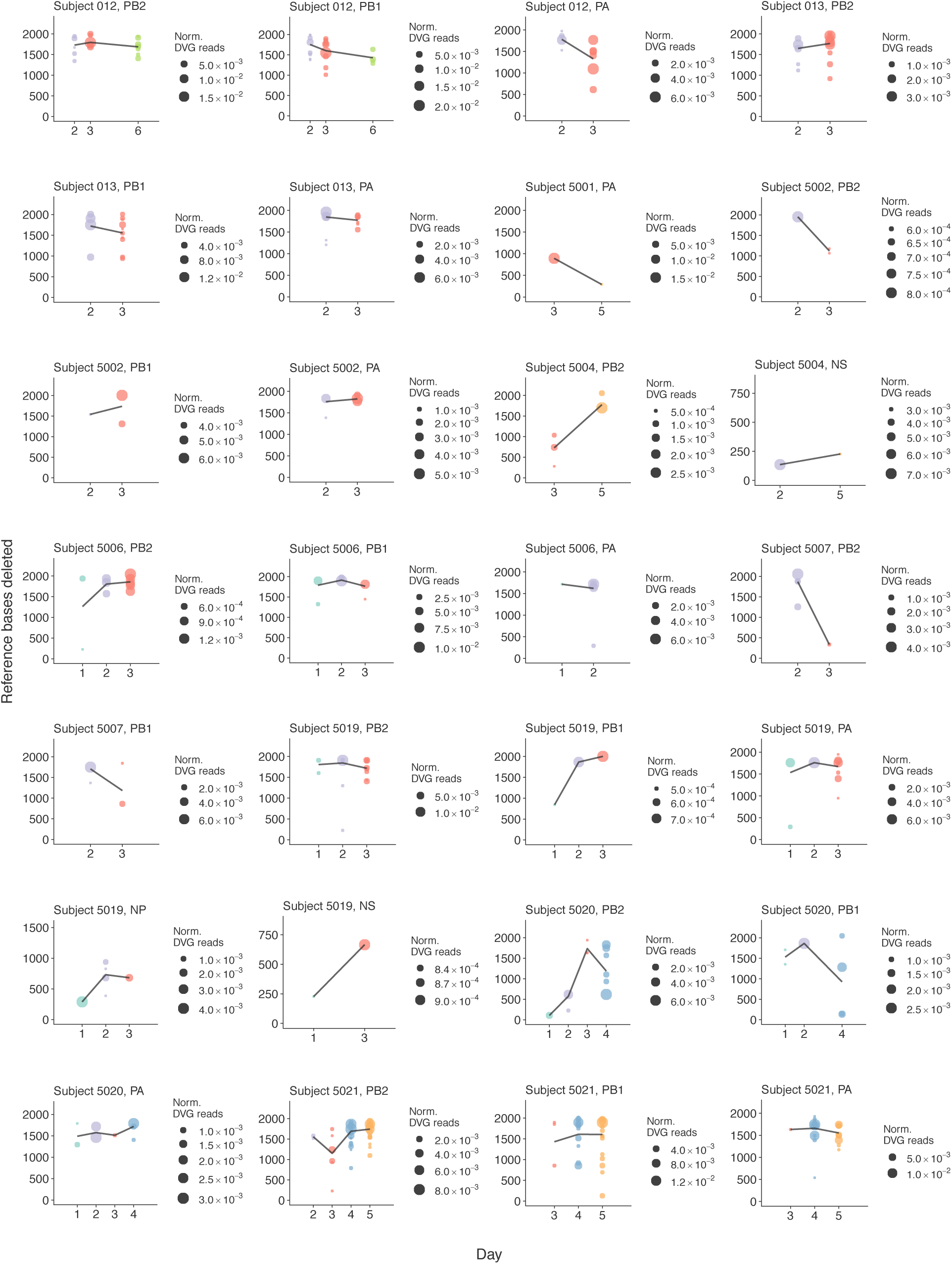
Dot plots representing the number of reference bases deleted in the observed defective viral genomes (DVG) species with each gene for subjects with DVGs observed on multiple days. Dot size is dependent on the number of reads supporting a given DVG species, normalized by the number of reads mapped to that gene segment. Color represents sampling day. Trend lines connect the mean number of reference bases deleted on each given day, weighted by the normalized number of supporting reads.

**TABLE S1.** Viral load and symptom scores Summary statistics for peak viral titer (log10(TCID50/mL), time to peak viral titer (hours), duration of infection, peak Modified Jackson symptom score, time to peak symptom score (hours), duration of symptoms (hours), and cumulative symptom score. Mean and population standard deviation are presented for all subjects, those in the early (treatment with oseltamivir on the evening of the first day post challenge) cohort, and those in the standard (treatment with oseltamivir on the evening of the fifth day post challenge) cohort. P-values comparing the early and standard treatment cohorts resultant from Mann-Whitney *U* tests are shown at right.

|  | Cohort |  |  | p-value |
| --- | --- | --- | --- | --- |
|  | All (mean [sd]) | Standard (mean [sd]) | Early (mean [sd]) |  |
| Peak viral titer (log<sub>10</sub>(TCID<sub>50</sub>/mL) | 4.46 [1.15] | 4.55 [0.84] | 4.34 [1.48] | 0.922 |
| Time to peak viral titer (hours) | 57.85 [25.75] | 64.95 [27.01] | 47.71 [19.82] | 0.080 |
| Duration of infection (hours) | 99.71 [38.57] | 117.80 [37.23] | 73.86 [22.32] | 0.035 |
| Peak SS | 6.47 [4.75] | 6.10 [4.25] | 7.00 [5.35] | 0.922 |
| Time to peak SS (hours) | 46.65 [22.18] | 59.30 [17.57] | 28.57 [14.09] | 0.003 |
| Duration of symptoms (hours) | 113.06 [36.98] | 134.20 [17.21] | 82.86 [36.71] | 0.005 |
| Cumulative SS | 39.88 [42.94] | 43.30 [49.99] | 35.00 [29.43] | 0.887 |

**TABLE S2.** Clinical data by subject Clinical data, including inoculum dose (log_10_(TCID_50_/mL)), treatment cohort, peak viral titer (log_10_(TCID_50_/mL)), time to peak viral titer (hours), duration of infection (hours), peak Modified Jackson symptom score, time to peak symptom score (hours), and cumulative symptom score, by subject

| Subject | Inoculum<br>(TCID<sub>50</sub>/ml) | Treatment<br>cohort | Peak viral<br>titer<br>(TCID<sub>50</sub>/ml) | Time to peak<br>viral titer<br>(hours) | Duration of<br>infection<br>(hours) | Peak<br>symptom<br>score | Time to peak<br>symptom score<br>(hours) | Cumulative<br>symptom<br>score |
| --- | --- | --- | --- | --- | --- | --- | --- | --- |
| 001 | 6.41 | Standard | 4.25 | 24.0 | 48.0 | 9 | 45.0 | 40 |
| 006 | 5.25 | Standard | 5.00 | 48.0 | 168.0 | 7 | 36.0 | 55 |
| 008 | 5.25 | Standard | 4.75 | 48.0 | 74.0 | 10 | 60.0 | 60 |
| 010 | 4.41 | Standard | 3.75 | 74.0 | 120.0 | 5 | 93.0 | 22 |
| 012 | 4.41 | Standard | 5.01 | 48.0 | 168.0 | 4 | 69.0 | 18 |
| 013 | 3.08 | Standard | 5.50 | 74.0 | 96.0 | 2 | 69.0 | 12 |
| 015 | 3.08 | Standard | 4.50 | 120.0 | 144.0 | 2 | 45.0 | 9 |
| 5001 | 5.50 | Early | 2.75 | 95.0 | 124.5 | 2 | 0.0 | 11 |
| 5002 | 5.50 | Early | 5.50 | 42.0 | 70.0 | 14 | 40.0 | 63 |
| 5004 | 5.50 | Standard | 2.50 | 76.5 | 118.0 | 3 | 56.0 | 23 |
| 5006 | 5.50 | Early | 6.25 | 42.0 | 76.5 | 8 | 40.0 | 45 |
| 5007 | 5.50 | Early | 4.00 | 42.0 | 70.0 | 2 | 32.0 | 6 |
| 5017 | 5.50 | Early | 1.75 | 42.0 | 53.0 | 1 | 16.0 | 1 |
| 5018 | 5.50 | Early | 5.00 | 29.0 | 53.0 | 7 | 40.0 | 33 |
| 5019 | 5.50 | Early | 5.12 | 42.0 | 70.0 | 15 | 32.0 | 86 |
| 5020 | 5.50 | Standard | 5.00 | 42.0 | 100.5 | 16 | 40.0 | 184 |
| 5021 | 5.50 | Standard | 5.27 | 95.0 | 141.5 | 3 | 80.0 | 10 |

**Table S3.**
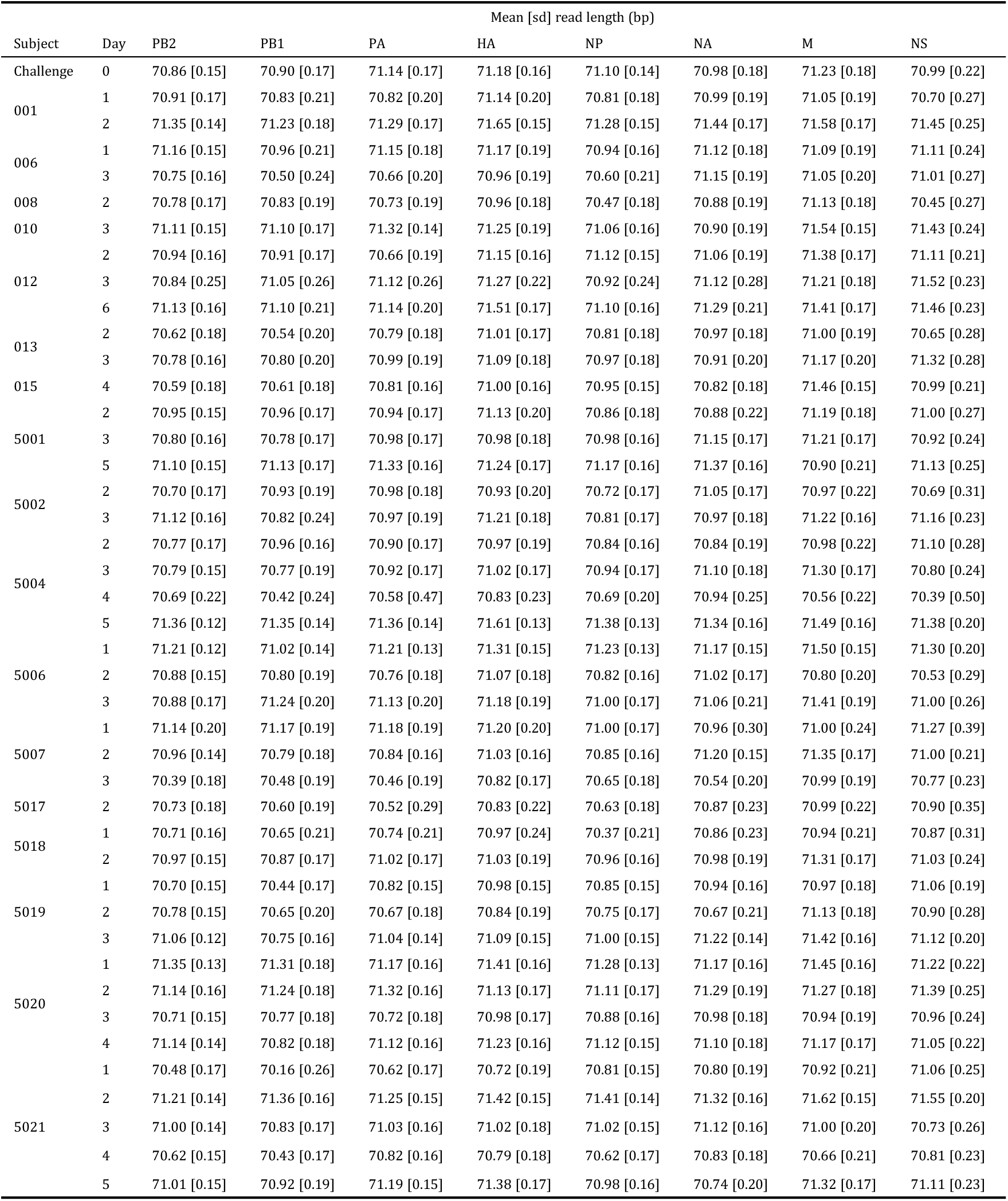
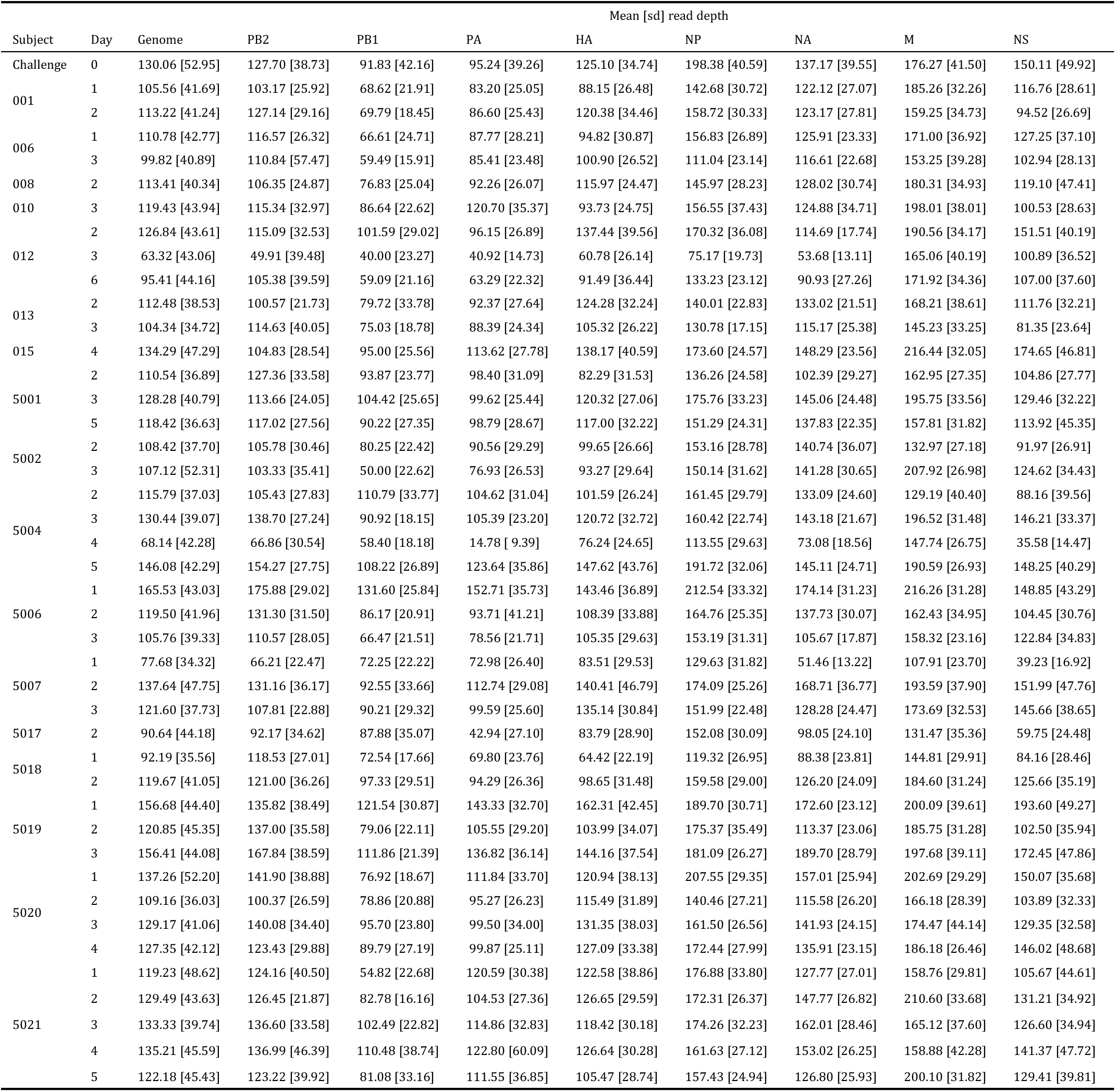
Sequencing data, including mean [population standard deviation (sd)] read length (bp) and mean [sd] coverage (reads), for all samples. Data are calculated following read trimming (removal of Illumina adapters, leading or trailing bases with quality <3, portions of reads where the average quality per-base in 4-base wide sliding windows was <15 and reads with <50 bases) (46), alignment to the reference genome, realignment to a consensus sequence (47), and removal of PCR duplicates (49). Technical sequencing replicates were processed separately and summary statistics calculated from the combination of the two samples.

**TABLE S4.** DVG read support Normalized DVG read count for all subjects for each gene segment. Values represent the proportion of the reads mapped to a given gene which support DVG species present in both technical replicates. Blank cells represent instances where no DVG reads were observed. Subjects highlighted in grey represent those in the early treatment (oseltamivir on the first day post challenge) treatment cohort.

| Subject | Day | Normalized DVG reads |  |  |  |  |  |  |  |
| --- | --- | --- | --- | --- | --- | --- | --- | --- | --- |
|  |  | PB2 | PB1 | PA | HA | NP | NA | M | NS |
| Challenge | 0 |  |  |  |  | 0.0014 | 0.0014 |  | 0.0021 |
| 1001 | 1 |  |  |  |  |  |  |  | 0.0013 |
|  | 2 | 0.0007 | 0.0026 | 0.0015 |  |  |  |  |  |
| 1006 | 1 |  |  |  |  | 0.0031 |  |  |  |
|  | 3 | 0.0529 | 0.0259 | 0.0154 |  |  | 0.0008 | 0.0009 |  |
| 1008 | 2 | 0.0006 | 0.0023 |  | 0.0007 |  |  |  |  |
| 1010 | 3 | 0.0099 | 0.0200 | 0.0641 | 0.0098 |  | 0.0030 |  |  |
| 1012 | 2 | 0.0072 | 0.0131 | 0.0071 |  |  |  |  |  |
|  | 3 | 0.0351 | 0.0452 | 0.0204 | 0.0013 |  |  |  |  |
|  | 6 | 0.0179 | 0.0096 |  |  |  | 0.0047 |  |  |
| 1013 | 2 | 0.0068 | 0.0365 | 0.0141 |  |  | 0.0007 |  |  |
|  | 3 | 0.0168 | 0.0181 | 0.0074 | 0.0008 |  | 0.0008 |  |  |
| 1015 | 4 |  | 0.0019 | 0.0016 |  |  |  |  | 0.0009 |
| 5001 | 3 |  |  | 0.0174 |  |  |  |  |  |
|  | 5 | 0.0005 |  | 0.0006 | 0.0007 |  | 0.0031 | 0.0026 | 0.0069 |
| 5002 | 2 | 0.0008 | 0.0033 | 0.0042 |  | 0.0011 |  |  |  |
|  | 3 | 0.0012 | 0.0106 | 0.0106 |  |  |  |  |  |
| 5004 | 2 |  |  | 0.0006 |  |  |  |  | 0.0072 |
|  | 3 | 0.0021 |  |  |  |  |  | 0.0007 |  |
|  | 4 |  |  |  |  |  | 0.0013 |  |  |
|  | 5 | 0.0038 | 0.0034 |  |  |  |  |  | 0.0028 |
| 5006 | 1 | 0.0009 | 0.0072 | 0.0004 |  |  |  |  |  |
|  | 2 | 0.0025 | 0.0132 | 0.0127 | 0.0015 |  |  |  | 0.0023 |
|  | 3 | 0.0055 | 0.0077 |  |  |  |  |  |  |
| 5007 | 2 | 0.0082 | 0.0083 | 0.0036 |  |  |  |  |  |
|  | 3 | 0.0006 | 0.0029 |  |  |  | 0.0018 | 0.0039 |  |
| 5018 | 1 |  |  |  |  | 0.0018 |  |  |  |
|  | 2 | 0.0044 | 0.0045 |  | 0.0076 |  |  |  |  |
| 5019 | 1 | 0.0020 | 0.0005 | 0.0057 |  | 0.0042 | 0.0023 |  | 0.0008 |
|  | 2 | 0.0157 | 0.0008 | 0.0077 |  | 0.0035 |  |  |  |
|  | 3 | 0.0132 | 0.0008 | 0.0129 |  | 0.0017 |  |  | 0.0009 |
| 5020 | 1 | 0.0021 | 0.0016 | 0.0014 |  |  |  |  |  |
|  | 2 | 0.0051 | 0.0027 | 0.0053 |  |  |  |  |  |
|  | 3 | 0.0013 |  | 0.0006 |  |  |  |  |  |
|  | 4 | 0.0170 | 0.0053 | 0.0044 |  |  | 0.0010 |  |  |
| 5021 | 2 | 0.0012 |  |  |  |  |  |  |  |
|  | 3 | 0.0062 | 0.0021 | 0.0006 |  |  |  |  |  |
|  | 4 | 0.0285 | 0.0351 | 0.0440 |  |  | 0.0057 |  |  |
|  | 5 | 0.0247 | 0.0392 | 0.0233 |  |  |  | 0.0007 | 0.0055 |

**TABLE S5.** DVG junction sites Summary statistics (mean [population standard deviation (sd)], weighted by the normalized DVG read count) for the 3’ and 5’ junction sites (relative to the reference) of DVGs observed in the PB2, PB1, and PA genes as well as the mean [sd] number of reference bases deleted.

|  | Reference<br>length (nt) | 3' junction site |  |  | 5' junction site |  |  | Mean [sd]<br>bases deleted<br>(nt) |
| --- | --- | --- | --- | --- | --- | --- | --- | --- |
|  |  | Mean [sd]<br>(nt) | Minimum<br>(nt) | Maximum<br>(nt) | Mean [sd]<br>(nt) | Minimum<br>(nt) | Maximum<br>(nt) |  |
| PB2 | 2310 | 325 [295] | 84 | 1814 | 1971 [236] | 398 | 2177 | 1646 [392] |
| PB1 | 2312 | 304 [259] | 43 | 2113 | 1929 [225] | 187 | 2224 | 1625 [397] |
| PA | 2192 | 256 [180] | 66 | 1455 | 1849 [204] | 243 | 2074 | 1593 [332] |

**TABLE S6.** DVG species observed in multiple samples Defective viral genome (DVG) species observed in multiple samples. DVGs are organized by gene segment. DVG IDs represent the first_last deleted reference based as identified using the “jI” SAM tag.

| Gene | DVG ID | Samples |  |  |  |
| --- | --- | --- | --- | --- | --- |
| PB2 | 93_1983 | 013, Day 2 | 013, Day 3 |  |  |
| PB2 | 154_2069 | 5019, Day 2 | 5019, Day 3 |  |  |
| PB2 | 170_2035 | 5021, Day 4 | 5021, Day 5 |  |  |
| PB2 | 171_2087 | 010, Day 3 | 5007, Day 2 |  |  |
| PB2 | 172_2079 | 5019, Day 1 | 5019, Day 2 | 5019, Day 3 |  |
| PB2 | 187_2125 | 5006, Day 1 | 5006, Day 2 | 5006, Day 3 |  |
| PB2 | 209_1983 | 010, Day 3 | 5006, Day 3 |  |  |
| PB2 | 211_1956 | 5021, Day 3 | 5021, Day 4 | 5021, Day 5 |  |
| PB2 | 241_1982 | 013, Day 2 | 013, Day 3 |  |  |
| PB2 | 249_2005 | 5021, Day 4 | 5021, Day 5 |  |  |
| PB2 | 356_1937 | 5021, Day 2 | 5021, Day 3 | 5021, Day 4 | 5021, Day 5 |
| PB2 | 386_1903 | 5021, Day 2 | 5021, Day 4 |  |  |
| PB2 | 476_703 | 5006, Day 1 | 5019, Day 2 | 5020, Day 2 | 5021, Day 3 |
| PB2 | 505_1743 | 5021, Day 3 | 5021, Day 4 |  |  |
| PB2 | 523_1789 | 013, Day 2 | 013, Day 3 |  |  |
| PB2 | 1482_2101 | 5020, Day 2 | 5020, Day 4 |  |  |
| PB1 | 43_1978 | 5021, Day 4 | 5021, Day 5 |  |  |
| PB1 | 90_2073 | 5021, Day 4 | 5021, Day 5 |  |  |
| PB1 | 131_2028 | 5006, Day 1 | 5006, Day 2 |  |  |
| PB1 | 142_2150 | 013, Day 2 | 013, Day 3 |  |  |
| PB1 | 179_2076 | 5021, Day 3 | 5021, Day 4 | 5021, Day 5 |  |
| PB1 | 183_1994 | 012, Day 2 | 012, Day 3 |  |  |
| PB1 | 183_2092 | 013, Day 2 | 013, Day 3 |  |  |
| PB1 | 226_1924 | 010, Day 3 | 5021, Day 5 |  |  |
| PB1 | 231_1746 | 5021, Day 4 | 5021, Day 5 |  |  |
| PB1 | 232_1986 | 013, Day 2 | 013, Day 3 |  |  |
| PB1 | 313_1863 | 012, Day 2 | 012, Day 3 |  |  |
| PB1 | 631_1603 | 013, Day 2 | 013, Day 3 |  |  |
| PB1 | 689_1544 | 5021, Day 3 | 5021, Day 4 | 5021, Day 5 |  |
| PA | 102_1936 | 5002, Day 2 | 5002, Day 3 |  |  |
| PA | 124_1891 | 012, Day 2 | 012, Day 3 |  |  |
| PA | 153_1941 | 5020, Day 1 | 5020, Day 4 |  |  |
| PA | 153_2015 | 013, Day 2 | 013, Day 3 |  |  |
| PA | 160_1903 | 5021, Day 4 | 5021, Day 5 |  |  |
| PA | 175_1892 | 5020, Day 2 | 5020, Day 4 |  |  |
| PA | 175_1935 | 5019, Day 1 | 5019, Day 2 | 5019, Day 3 |  |
| PA | 197_1916 | 5021, Day 4 | 5021, Day 5 |  |  |
| PA | 229_1944 | 5006, Day 1 | 5006, Day 2 |  |  |
| PA | 263_1968 | 5021, Day 4 | 5021, Day 5 |  |  |
| PA | 328_1719 | 5021, Day 4 | 5021, Day 5 |  |  |
| PA | 331_1824 | 5021, Day 4 | 5021, Day 5 |  |  |
| PA | 476_767 | 5001, Day 5 | 5004, Day 2 | 5006, Day 2 | 5019, Day 1 |
| NP | 229_518 | 006, Day 1 | 5018, Day 1 | 5019, Day 1 |  |
| NP | 696_1378 | 5002, Day 2 | 5019, Day 2 | 5019, Day 3 | Challenge, Day 0 |
| NA | 421_803 | 006, Day 3 | 013, Day 2 |  |  |
| NA | 633_737 | 010, Day 3 | 5019, Day 1 | 5021, Day 4 |  |
| NS | 291_518 | 001, Day 1 | 5004, Day 5 | 5019, Day 1 | 5021, Day 5 |

